## Supplemental Data for "The small acid-soluble proteins of *Clostridioides difficile* regulate sporulation in a SpoIVB2-dependent manner"

**Supplement Figure 1. EMS mutagenesis strategy.** EMS was added to logarithmically growing cultures and incubated for 3 hours. The resulting cells were spread on sporulation medium (70:30) and incubated for 5 days. Subsequently, the growth was harvested, purified, and then then heat shocked at 65 °C for 1 hour. After heat treatment, the samples were plated onto 70:30 and again incubated for 5 days. This enrichment process was repeated 3 times before individual colonies were isolated, phenotypes confirmed, and DNA sent for whole genome re-sequencing. The figure was generated using BioRender.com.

**Supplement Figure 2. Spore yield is reduced with expression of *spoIVB2* in EMS strains*.*** Spore yield of the indicated strain was determined as described in Figure 1. pEV indicates an empty vector. A) Strains isolated during EMS. B) Clean strains containing generated *spoIVB2* alleles. All data represents the average of three independent experiments. Statistical analysis by one way ANOVA with Šídák's multiple comparison test. * P<0.05, ** P<0.01, *** P<0.001, **** P<0.0001. B) **a** P<0.0001 in comparison to *C. difficile* Δ*sspA* Δ*sspB*.

**Supplement Figure 3. qPCR of mutant strains: Part 1*.*** Strains were grown on sporulation medium for 11 hours before RNA extraction. qPCR was performed using SYBR green. Transcripts for the following genes were determined: A) *sspA*, B) *sspB*, C) *spoIVA*, D) *sleC*, E) *pdaA*, F) *spoVT.* Fold change from R20291 was determined with the ΔΔCT method using *rpoA* transcripts as the internal control. All data represents the average of five independent experiments. Statistical analysis by one way ANOVA with Dunnett's multiple comparison test with the mutant strains compared to wild type. * P<0.05, ** P<0.01, *** P<0.001, **** P<0.0001.

**Supplement Figure 4. qPCR of mutant strains: Part 2*.*** Strains were grown on sporulation medium for 11 hours before RNA extraction. qPCR was performed using SYBR green. Transcripts for the following genes were determined: A) *spoIVB*, B) *spoIVB2*, C) *spoIIP.* Fold change from R20291 was determined with the ΔΔCT method using *rpoA* transcripts as the internal control. All data represents the average of five independent experiments. Statistical analysis by one way ANOVA with Dunnett's multiple comparison test with the mutant strains compared to wild type. * P<0.05, ** P<0.01, *** P<0.001, **** P<0.0001.

**Supplement Figure 5. qPCR of mutant strains: Part 3*.*** Strains were grown on sporulation medium for 11 hours before RNA extraction. qPCR was performed using SYBR green. Transcripts for the following genes were determined: A) *dpaA*, B) *spoVAC*, C) *spoVAD*, D) *spoVAE.* Fold change from R20291 was determined with the ΔΔCT method using *rpoA* transcripts as the internal control. All data represents the average of five independent experiments. Statistical analysis by one way ANOVA with Dunnett's multiple comparison test with the mutant strains compared to wild type. * P<0.05, ** P<0.01, *** P<0.001, **** P<0.0001.

**Supplement Table 3. Mutations found within suppressor strains.** The mutations identified in EMS treated suppressor strains are sorted by position within the genome and color-coded based on the isolate containing the mutation.

Supplement Table 1. Primers used in this study.

| Primer Name | Sequence |
| --- | --- |
| 5'sspA_MTL | ttatcaggaaacagctatgaccgcggccgcttagatgaggaaaaactggataa |

| | |
|---|---|
| 3'sspA_up | ttatttataactatctgttgctttttccaggttgattaccttccttctgttta |
| 5'sspA_down | aataaattaaacagaaggaaggtaatcaacctggaaaaagcaacagatagt |
| 3' sspA_xylR | tgcaggcttcttatttttatgctagctcgagctattgaacttggaaatgagag |
| CRISPR_sspA_165 | gtgtgctataattaaactgtaaaacgcgtgactaaaaaattagttgaaagttttagagctagaaatagcaagttaaaataaggctagtccgttatcaacttgaaaaagtggcaccgagtcggtgctttttttctatggagaaatctagatcagcatgatgtctgactagacgcgtaagctctgcaactatttttagat |
| 5'traJ | gcgaggaagcggaagagcgcccaatacgcagggccccctgcttcggggtca |
| 3'traJ | aatttatctacaattttttatcctgcagggggcccgatcggtcttgccttg |
| CRISPR_sspA_135 | taattaaactgtaaaggtaccagagaaaatggttatgtagggttttagagctagaaatagcaagttaaaataaggctagtccgttatcaacttgaaaaagtggcaccgagtcggtgctttttttctatggagaaatctagatcagcatgatgtctgactagacgcgtaagctctgcaactat |
| 5' sspB UP | attttttatcaggaaacagctatgaccgcggccgcttttaaaatatcatccatattat |
| 3' sspB UP | tgtcaaaatttactatttattttccagccacctcaaataattagtttatgatg |
| 5' sspB DN | tgttagacatcataaactaattatttgaggtggctggaaaataaatagta |
| 3' sspB_xylR | atgcaggcttcttatttttatgctagctcgagatacttgtctattttttcagtaa |
| CRISPR_sspB_144 | aattaaactgtaaaggtaccagagaaaatggatatgttgggttttagagctagaaatagcaagttaaaataaggctagtccgttatcaacttgaaaaagtggcaccgagtcggtgctttttttctatggagaaatctagatcagcatgatgtctgactagacgcgtaagctctgcaacta |
| 5' CDR20291_0714 UP | ttatcaggaaacagctatgaccgcggccgcccttgatgccctgggtctatac |
| 3' CDR20291_0714 UP | cttatttatttatattatttacaacatccgttgcataaaacacctctttct |
| 5' CDR20291_0714 DN | ttaataagaaagaggtgttttatgcaacggatgttgtaaataatataaataa |
| 3' CDR20291_0714 DN | gcaggcttcttatttttatgctagctcgagggaacagcattaggaagtcc |
| CDR20291_0714 gRNA 3 | gcattcaaggagggggtaccgtatctattttattaaataggttttagagctagaaatagc |
| 3' gRNA_change | ccatctaaaaatagttgcagagcttacgcgtctagtcagacatcatgctgatctag |
| 5' tn916.traJ | tctgcagattacctaataatttatctacattccctttagcctgcttcggggtcattat |
| 5'Tn916ori_gibson | cggaagagcgcccaatacgcagggccctaacatcttctatttttcccaaatc |
| 3' tn916.traJ | cgaaaaaatcgctataatgaccccgaagcaggctaaagggaatgtagataaattattag |
| 5' CDR20291_0714 | tttttatcaggaaacagctatgaccgcggccgctgctatctccttttccttg |
| 3' CDR20291_0714 | gtgccaagcttgcatgtctgcaggcctcgagttacaacatccatttaataaatac |
| 3' 0714_S301A | ttattatcttgaactattggagtacctgccataccttgtacaataccacc |

| | |
|---|---|
| 5' 0714_S301A | ctcaaactggtggtattgtacaaggtatggcaggtactccaatagttcaag |
| 3' sspA.pJS116 | tgccaagcttgcatgtctgcaggcctcgagctatctgttgctttttccag |
| 3' sspAsspB | ggaactgataatatggatgatattttaaaactatctgttgctttttccagccatttg |
| 5' sspAsspB | caaatggctggaaaaagcaacagatagttttaaaatatcatccatattat |
| 3'sspBpJS116 | cagtgccaagcttgcatgtctgcaggcctcgagttattttccagccatttgtc |
| 5' rpoA | taaaggtagaggttatgtttctgct |
| 3' rpoA | tttgaccaactcttgtgttttcc |
| 5' sspA_qPCR | caaaagaggctttaaaccaaatgaa |
| 3' sspA_qPCR | attttctcttgcagtaaggtttcctt |
| 5' sspB_qPCR | aacagaacagtagttccagaagcaaa |
| 3' sspB_qPCR | caacatatccattttctctagctgttaag |
| 5'sleC_qPCR | ttgaagcaagacaaggagttccc |
| 3'sleC_qPCR | cgaaaccagtaggaggaggtaatgg |
| 5' spoVT_qPCR | agagaaggagaccctttagagat |
| 3' spoVT_qPCR | ctgttatcaacactccatatcctagt |
| 5' pdaA_qPCR | tggtaaacagccatcacctataa |
| 3' pdaA_qPCR | tccactttcatatccagcatca |
| 5'spoIVA_qPCR | ggatagaacaagagatgagataccc |
| 3'spoIVA_qPCR | ctgctgccttttcaaatgtc |
| 5' spoIVB_qPCR_1 | ttcagagctaggtataagtggtaat |
| 3' spoIVB_qPCR_1 | tgcggtcttccaacttctatt |
| 5' spoIVB2_qPCR_1 | agctcaaactggtggtattgt |
| 3' spoIVB2_qPCR_1 | catgtgacactgctccgatta |
| 5' spoIIP_qPCR_1 | catactcatggatgtgagacttattcaa |
| 3' spoIIP_qPCR_1 | accccatcctttgctatctaaagc |
| 5' dpaA_qPCR | actgttattgggggagacctgc |
| 3' dpaA_qPCR | ggaacttttggatagtgcttcagc |

| | |
|---|---|
| 5' spoVAC_qPCR | agctggagctggttctataattcc |
| 3' spoVAC_qPCR | catagccttctcttttatactccattgc |
| 5' spoVAD_qPCR | tgacagctcagtggacagttacag |
| 3' spoVAD_qPCR | tttggaccgtctccattagga |
| 5' spoVAE_qPCR | gtttaatagcccaagtaatgatggatt |
| 3' spoVAE_qPCR | acaccagttgttacatacgttaccataa |
| 3' luciferase_ssrA_pHN149 | aagcttgcatgtctgcaggcctcgagtcatcatgcagcaagtgcataattttcatcattagctgctagaatt tcttcaaaaagtctat |
| 3' luciferase_pHN149 | gccaagcttgcatgtctgcaggcctcgagtcatcatagaatttcttcaaaaag |
| 3' PsspA_BS49 | gttgttgctgttacctgagttattgttagccatgttgattaccttccttctgt |
| 5' sspA_BS49 | acacaaaataaattaaacagaaggaaggtaatcaacatggctaacaataactcagg |
| 3' sspA_BS49 | ccagtgccaagcttgcatgtctgcaggcctcgagttagaattgtcctccgcc |
| 3' spoIVB2 F36F | attttgtgcataaattaaattatttgagaagaaatataataaaaataaaaatgttaaaac |
| 5' spoIVB2 F36F | tacaattgttttaacatttttatttttattatatttcttctcaaataatttaatttatgc |
| 3' spoIVB2 F37.UUA | aattttgtgcataaattaaattatttgataaaaaatataataaaaataaaaatgttaaa |
| 5' spoIVB2 F37.UUA | caattgttttaacatttttatttttattatattttttatcaaataatttaatttatgcac |
| 3' spoIVB2 F37.UUG | aattttgtgcataaattaaattatttgacaaaaaatataataaaaataaaaatgttaaa |
| 5' spoIVB2 F37.UUG | caattgttttaacatttttatttttattatattttttgtcaaataatttaatttatgcac |
| 5’sacB_UP | ttatcaggaaacagctatgaccgcggccgcgtcgactagttctttaggcccg |
| sacB_3’_XhoI | aagcttgcatgtctgcaggcctcgagttatttgttaactgttaattgtccttgttcaagg |
| 3’ PsspA_spoIVB2 | ttaaagtattatttttaaaatgaaaatttttaagttgcatgttgattaccttccttctg |
| 5’ spoIVB2_PsspA | acaaaataaattaaacagaaggaaggtaatcaacatgcaacttaaaaattttcatt |
| 5’ spoIVB.pHN149 | acaattttttatcaggaaacagctatgaccgcggccgctttattgtcttcgaatatac |
| 3’ PspoIVB_spoIVB2 | aagtattatttttaaaatgaaaatttttaagttgcatatatccatctactcctatgc |
| 5’ spoIVB2_PspoIVB | aatacataataatacagcataggagtagatggatatatgcaacttaaaaattttcat |
| 5’ PspoIVB2_pHN149 | attttttatcaggaaacagctatgaccgcggccgctatttatttttatgaaaactaagg |
| 3’ PspoIVB(100)_PspoIVB2 | cttaataagtgattttaatacataatatgaaaacacctctttcttatta |
| 5’ PspoIVB2_PspoIVB | ctattattaaaataatttaataagaaagaggtgttttcatattatgtattaaaaatcact |

| | |
|---|---|
| 3' PsspA_PspoIVB2 | ttaattcaggtaatttttagcaattaaaacctgaaaacacctctttcttatta |
| 5' PsspA_PspoIVB2 | ttattaaaataatttaataagaaagaggtgttttcaggttttaattgctaaaaa |
| 3' spoIVB2_homol | tatttttaaaatgaaaatttttaagttgcataaaacacctctttcttattaaattat |
| 5' spoIVB2_gene_homol | ttaaaataatttaataagaaagaggtgttttatgcaacttaaaaattttcattttaa |
| 3' spoIVB2end_lrgBit | tcccaatcacccacaaaatcttcaagtgtaaacaccaacatccatttaataaatacacc |
| 5' lrgBit_spoIVB2end | ctgtaggttatggtgtatttattaaatggatgttggtgtttacacttgaagattttgtgg |
| 3' spoIVB2_bitLuc | ccaatcacccacaaaatcttcaagtgtaaacaccataaaacacctctttcttattaaatt |
| 5' bitLuc_PspoIVB2 | attattaaaataatttaataagaaagaggtgttttatggtgtttacacttgaagattttg |
| 5' spoIVB2_theo | ttttttatcaggaaacagctatgaccgcggccgcgatttgtcttcatttttatcttta |
| 3' spoIVB2_theo | gccagtgccaagcttgcatgtctgcaggcctcgagaaatatcaaagttattaatttgac |

CRISPR targeting sequence is in **bold.**

**Supplement Table 2. Strains and plasmids used in this study.**

| Strain | Description | Reference |
|---|---|---|
| *E. coli* DH5a | Cloning strain | [1] |
| *E. coli* HB101 pRK24 | Conjugal donor strain, $Amp^R$ | [2] |
| *E. coli* MB3436 | *recA*$^+$ *E. coli* strain | Gift from Dr. Michael Benedik |
| *B. subtilis* BS49 | *Tn*916 donor strain, $Tet^R$ | [3] |
| *C. difficile* R20291 | Wild type, ribotype 027 | [4] |
| *C. difficile* CD630Δ*erm* | Wild type, ribotype 012 | Gift from Dr. Daniel Paredes-Sabja (Texas A&M University) |
| *C. difficile* KNM10 | R20291 *spo0A* CRISPR-Cas9 mutant | [5] |
| *C. difficile* HNN03 | R20291 *sspA* CRISPR-Cas9 mutant | [6] |
| *C. difficile* HNN04 | R20291 *sspB* CRISPR-Cas9 mutant with an $sspA_{G52V}$ allele (called *sspB** throughout this manuscript) | [6] |
| *C. difficile* HNN05 | R20291 *sspA* and *sspB* CRISPR-Cas9 double mutant | [6] |
| *C. difficile* HNN17 | R20291 *sspB* CRISPR-Cas9 mutant | [6] |
| *C. difficile* HNN19 | EMS isolate from treatment of HNN04 | This study |

| | | |
|---|---|---|
| *C. difficile* HNN22 | EMS isolate from treatment of HNN04 | This study |
| *C. difficile* HNN26 | EMS isolate from treatment of HNN04 | This study |
| *C. difficile* HNN28 | EMS isolate from treatment of HNN04 | This study |
| *C. difficile* HNN32 | EMS isolate from treatment of HNN05 | This study |
| *C. difficile* HNN33 | EMS isolate from treatment of HNN05 | This study |
| *C. difficile* HNN35 | EMS isolate from treatment of HNN05 | This study |
| *C. difficile* HNN37 | EMS isolate from treatment of HNN05 | This study |
| *C. difficile* HNN38 | EMS isolate from treatment of HNN05 | This study |
| *C. difficile* HNN39 | EMS isolate from treatment of HNN05 | This study |
| *C. difficile* HNN40 | EMS isolate from treatment of HNN05 | This study |
| *C. difficile* HNN41 | EMS isolate from treatment of HNN05 | This study |
| *C. difficile* HNN43 | CD630Δ*erm sspB* CRISPR-Cas9 mutant | This study |
| *C. difficile* HNN45 | CD630Δ*erm sspA* CRISPR-Cas9 mutant | This study |
| *C. difficile* HNN46 | CD630Δ*erm sspA* and *sspB* CRISPR-Cas9 double mutant | This study |
| *C. difficile* HNN48 | EMS isolate from treatment of HNN05 | This study |
| *C. difficile* HNN49 | R20291_*0714* (*spoIVB2*) CRISPR-Cas9 mutant | This study |
| *C. difficile* HNN51 | EMS isolate from treatment of HNN05 | This study |
| *C. difficile* HNN57 | R20291 $spoIVB2_{F37F}$ | This study |
| *C. difficile* HNN60 | R20291 $spoIVB2_{A20T}$ | This study |
| *C. difficile* HNN64 | R20291 Δ*sspA* Δ*sspB* $spoIVB2_{A20T}$ | This study |
| *C. difficile* HNN73 | R20291 Δ*sspA* Δ*sspB* $spoIVB2_{F37F}$ | This study |
| **Plasmid** | **Description** | **Reference** |
| pMTL84151 | *E. coli* – *C. difficile* shuttle vector | [7] |
| pMTLYN4 | *traJ* containing plasmid | [8] |
| pJS116 | *B. subtilis* – *C. difficile* shuttle vector | [9] |
| pKM197 | CRISPR plasmid with *xylR* promoter driving *cas9* | [10] |
| pMB81 | BitLuc containing plasmid | [12] |

| | | |
|---|---|---|
| pJB09 | *cas*9 containing plasmid for the 2-plasmid CRISPR system | [13] |
| pJB14 | Targeting plasmid for the 2-plasmid CRISPR system | [13] |
| pJB94 | Theophylline allelic exchange base plasmid | [14] |
| pJB96 | pHN149 with *sacB* between *Not*I and *Xho*I cut sites, for easy selection of inserts | This study |
| pHN14 | R20291 *sspB* promoter region and gene | [6] |
| pHN30 | R20291 *sspA* and *sspB* complement | [6] |
| pHN120 | CD630Δ*erm sspA* targeted CRISPR vector, gRNA 1398 | This study |
| pHN121 | CD630Δ*erm sspB* targeted CRISPR vector, gRNA 1186 | This study |
| pHN122 | CDR20291_0714 promoter region and F37F allele | This study |
| pHN123 | CDR20291_0714 promoter region and A20T allele | This study |
| pHN127 | CDR20291_0714 promoter region and WT allele | This study |
| pHN131 | CD630Δ*erm sspA* targeted CRISPR vector with *TraJ oriT,* gRNA 165 | This study |
| pHN132 | CD630Δ*erm sspB* targeted CRISPR vector with *TraJ oriT*, gRNA 144 | This study |
| pHN138 | CD630Δ*erm sspA* targeted CRISPR vector with *TraJ oriT*, gRNA 135 | This study |
| pHN145 | CDR20291_0714 promoter region and S301A allele | This study |
| pHN146 | CDR20291_0714 promoter region and F37F, S301A allele | This study |
| pHN147 | CDR20291_0714 promoter region and A20T, S301A allele | This study |
| pHN149 | pMTL84151 based plasmid that also contains the *Tn916 oriT* (base plasmid that can be conjugated through *E. coli* or *B. subtilis* conjugal donors) | This study |
| pHN152 | CD630Δ*erm sspA* promoter region and gene | This study |
| pHN153 | CD630Δ*erm sspA* and *sspB* promoter region and gene | This study |
| pHN157 | CDR20291_0714 targeted CRISPR vector, gRNA 3 | This study |
| pHN176 | CD630Δ*erm sspB* promoter region and gene | This study |
| pHN208 | CDR20291_0714 promoter region and F36F | This study |
| pHN218 | CDR20291_0714 promoter region and F37L (UUA codon) | This study |
| pHN219 | CDR20291_0714 promoter region and F37L (UUG codon) | This study |
| pHN220 | *sspA* promoter region and *sspA* gene from *B. subtilis* BS49 | This study |
| pHN271 | *spoIVB2*A20T theophylline allelic exchange | This study |

| | | |
|---|---|---|
| pHN272 | *spoIVB2*F37F theophylline allelic exchange | This study |
| pHN312 | *sspA* promoter driving *spoIVB2* expression | This study |
| pHN329 | *spoIVB* promoter driving *spoIVB2* expression | This study |
| pHN330 | *spoIVB2* and *spoIVB* promoters driving *spoIVB2* expression | This study |
| pHN331 | *spoIVB2* and *sspA* promoters driving *spoIVB2* expression | This study |
| pHN335 | *spoIVB2* promoter with *spoIVB2* attached to *bitLuc* (luciferase) and tagged with *ssrA* | This study |
| pHN336 | *spoIVB2* promoter with *spoIVB2*$_{A20T}$ attached to *bitLuc* (luciferase) and tagged with *ssrA* | This study |
| pHN337 | *spoIVB2* promoter with *spoIVB2*$_{F37F}$ attached to *bitLuc* (luciferase) and tagged with *ssrA* | This study |
| pHN338 | *spoIVB2* promoter with *bitLuc* (luciferase) and tagged with *ssrA* | This study |
| pHN339 | *spoIVB2* promoter with *bitLuc* (luciferase) | This study |

1. Hanahan D. Studies on transformation of *Escherichia coli* with plasmids. J Mol Biol. 1983;166(4):557-80. Epub 1983/06/05. doi: 10.1016/s0022-2836(83)80284-8. PubMed PMID: 6345791.
2. Ma NJ, Moonan DW, Isaacs FJ. Precise manipulation of bacterial chromosomes by conjugative assembly genome engineering. Nature protocols. 2014;9(10):2285-300. doi: 10.1038/nprot.2014.081.
3. Bouillaut L, McBride SM, Sorg JA. Genetic Manipulation of *Clostridium difficile*. Current Protocols in Microbiology2011.
4. Stabler RA, He M, Dawson L, Martin M, Valiente E, Corton C, et al. Comparative genome and phenotypic analysis of *Clostridium difficile* 027 strains provides insight into the evolution of a hypervirulent bacterium. Genome Biol. 2009;10(9):R102. Epub 2009/09/29. doi: 10.1186/gb-2009-10-9-r102. PubMed PMID: 19781061; PubMed Central PMCID: PMCPMC2768977.
5. McAllister KN, Martinez Aguirre A, Sorg JA. The selenophosphate synthetase, *selD*, is important for *Clostridioides difficile* physiology. BioRxIV. 2021. doi: 10.1101/2021.01.06.425661.
6. Nerber HN, Sorg JA. The small acid-soluble proteins of *Clostridioides difficile* are important for UV resistance and serve as a check point for sporulation. PLoS Pathog. 2021;17(9):e1009516. Epub 2021/09/09. doi: 10.1371/journal.ppat.1009516. PubMed PMID: 34496003.
7. Heap JT, Pennington OJ, Cartman ST, Minton NP. A modular system for *Clostridium* shuttle plasmids. J Microbiol Methods. 2009;78(1):79-85. Epub 20090513. doi: 10.1016/j.mimet.2009.05.004. PubMed PMID: 19445976.
8. Ng YK, Ehsaan M, Philip S, Collery MM, Janoir C, Collignon A, et al. Expanding the repertoire of gene tools for precise manipulation of the *Clostridium difficile* genome: allelic exchange using *pyrE* alleles. PLoS One. 2013;8(2):e56051. Epub 2013/02/14. doi: 10.1371/journal.pone.0056051. PubMed PMID: 23405251; PubMed Central PMCID: PMCPMC3566075.

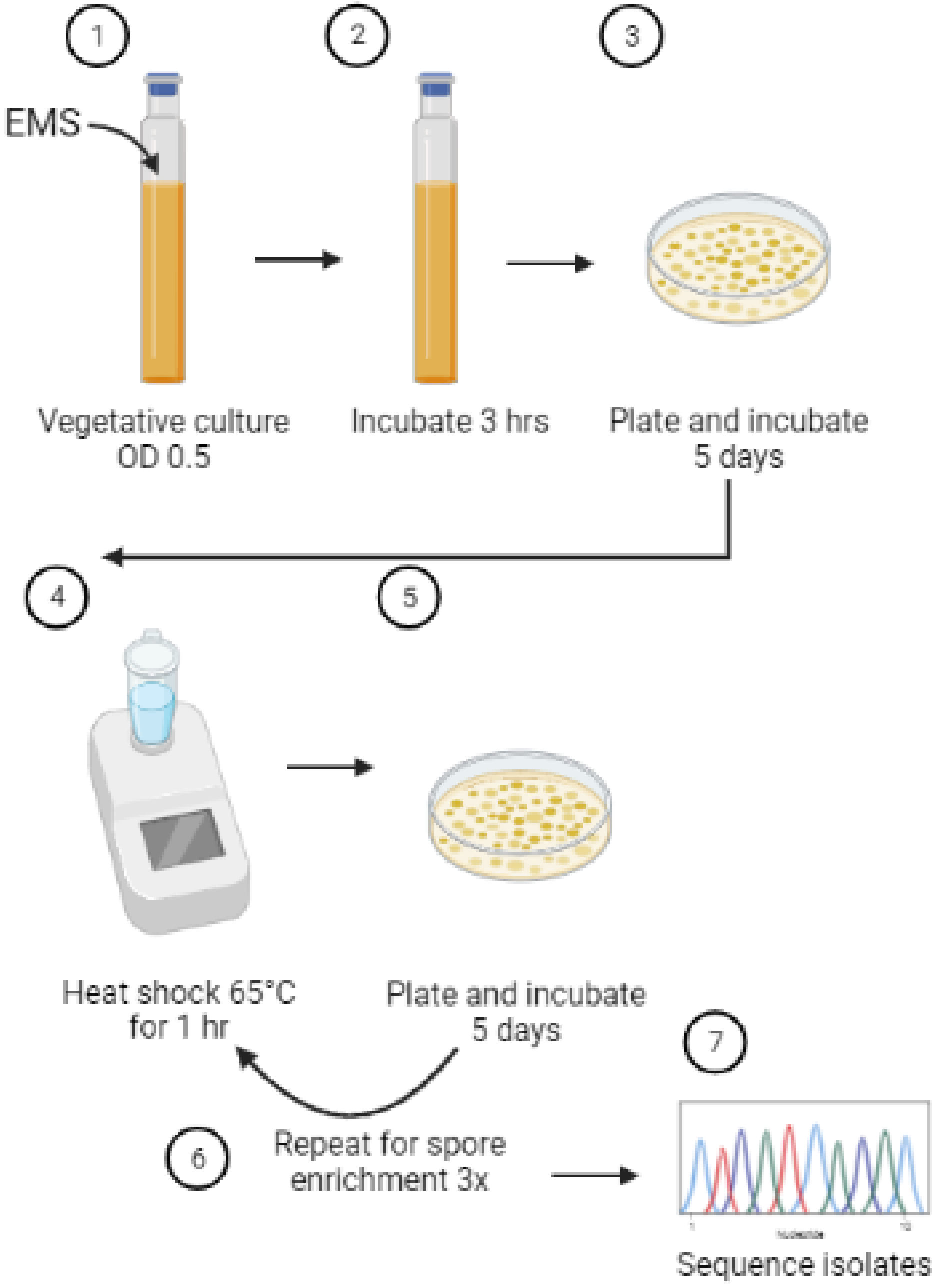
1
2
3
EMS
Vegetative culture
OD 0.5
Incubate 3 hrs
Plate and incubate
5 days
4
5
Heat shock 65°C
for 1 hr
Plate and incubate
5 days
7
6
Repeat for spore
enrichment 3x
Sequence isolates

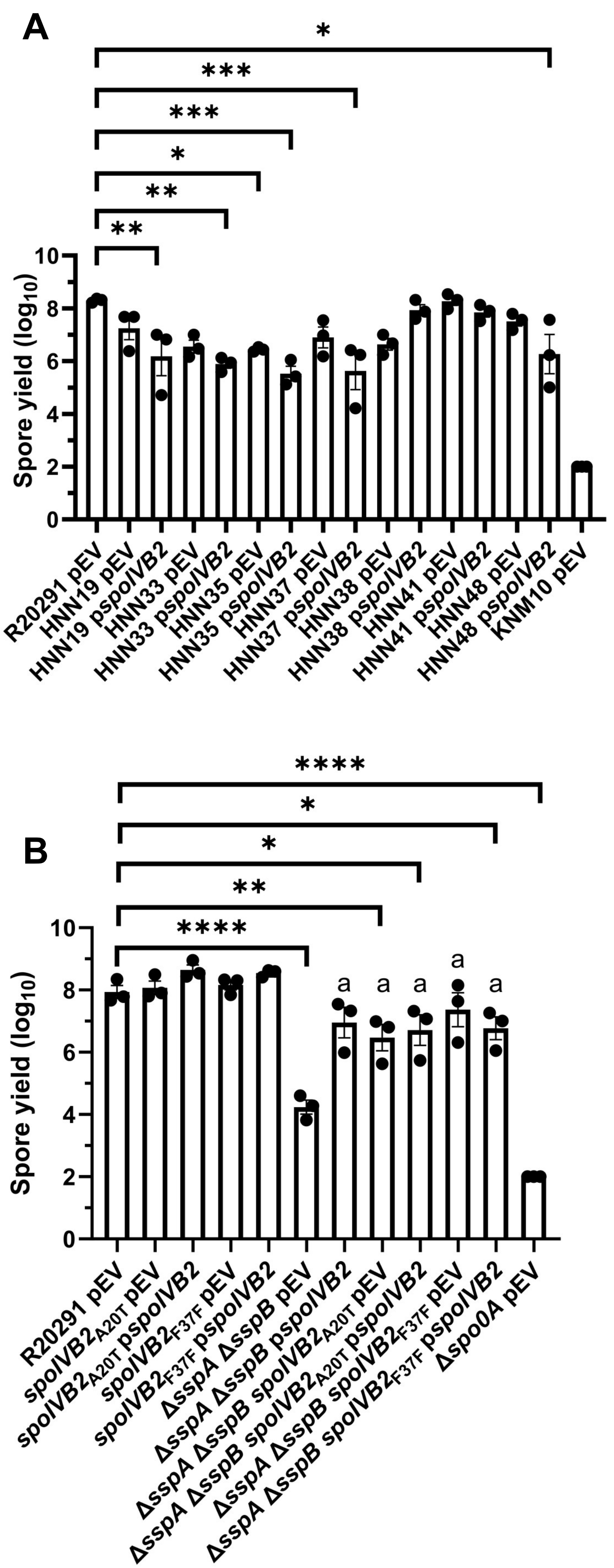
A
Spore yield (log10)
R20291 pEV
HNN19 pEV
HNN19 pspoIVB2
HNN33 pEV
HNN33 pspoIVB2
HNN35 pEV
HNN35 pspoIVB2
HNN37 pEV
HNN37 pspoIVB2
HNN38 pEV
HNN38 pspoIVB2
HNN41 pEV
HNN41 pspoIVB2
HNN48 pEV
HNN48 pspoIVB2
KNM10 pEV
B
Spore yield (log10)
R20291 pEV
spoIVB2A20T pEV
spoIVB2A20T pspoIVB2
spoIVB2F37F pEV
spoIVB2F37F pspoIVB2
ΔsspA ΔsspB pEV
ΔsspA ΔsspB pspoIVB2
ΔsspA ΔsspB spoIVB2A20T pEV
ΔsspA ΔsspB spoIVB2A20T pspoIVB2
ΔsspA ΔsspB spoIVB2F37F pEV
ΔsspA ΔsspB spoIVB2F37F pspoIVB2
Δspo0A pEV

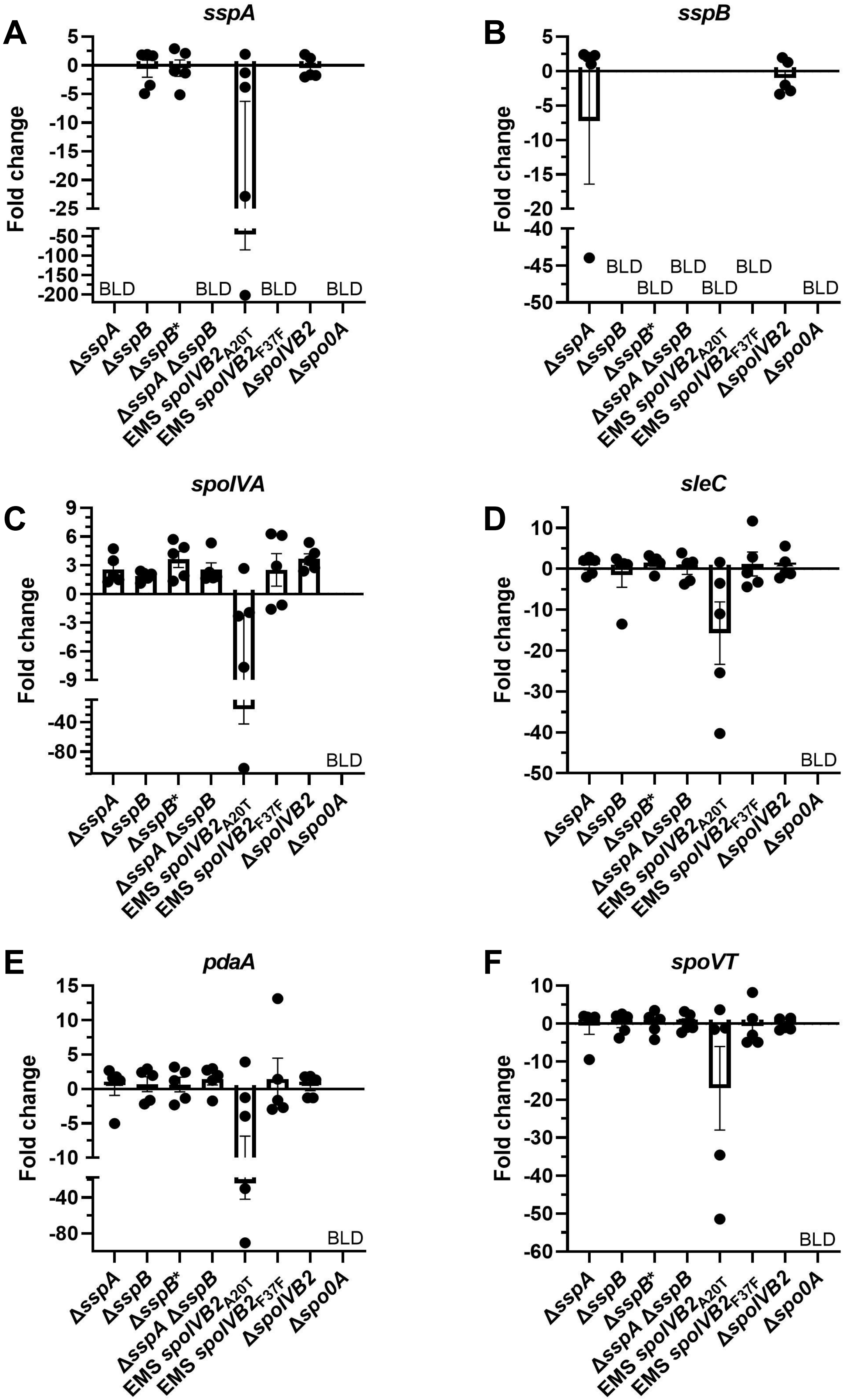

A
sspA
B
sspB
C
spoIVA
D
sleC
E
pdaA
F
spoVT
Fold change
BLD
ΔsspA
ΔsspB
ΔsspB*
ΔsspA ΔsspB
EMS spoIVB2A20T
EMS spoIVB2F37F
ΔspoIVB2
Δspo0A

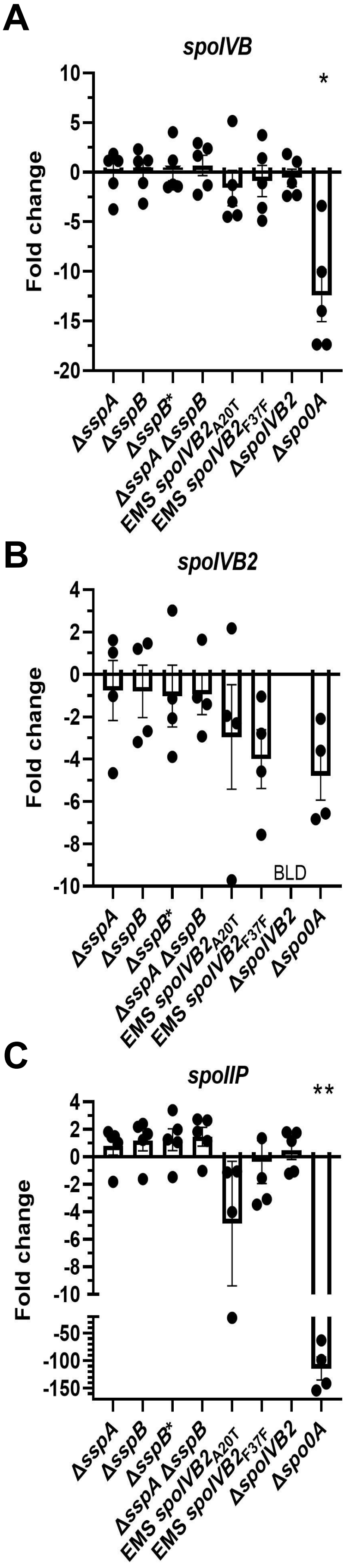

A
spoIVB
*
10
5
0
-5
-10
-15
-20
Fold change
ΔsspA
ΔsspB
ΔsspB*
ΔsspA ΔsspB
EMS spoIVB2A20T
EMS spoIVB2F37F
ΔspoIVB2
Δspo0A
B
spoIVB2
4
2
0
-2
-4
-6
-8
-10
Fold change
BLD
ΔsspA
ΔsspB
ΔsspB*
ΔsspA ΔsspB
EMS spoIVB2A20T
EMS spoIVB2F37F
ΔspoIVB2
Δspo0A
C
spoIIP
**
4
2
0
-2
-4
-6
-8
-10
-50
-100
-150
Fold change
ΔsspA
ΔsspB
ΔsspB*
ΔsspA ΔsspB
EMS spoIVB2A20T
EMS spoIVB2F37F
ΔspoIVB2
Δspo0A

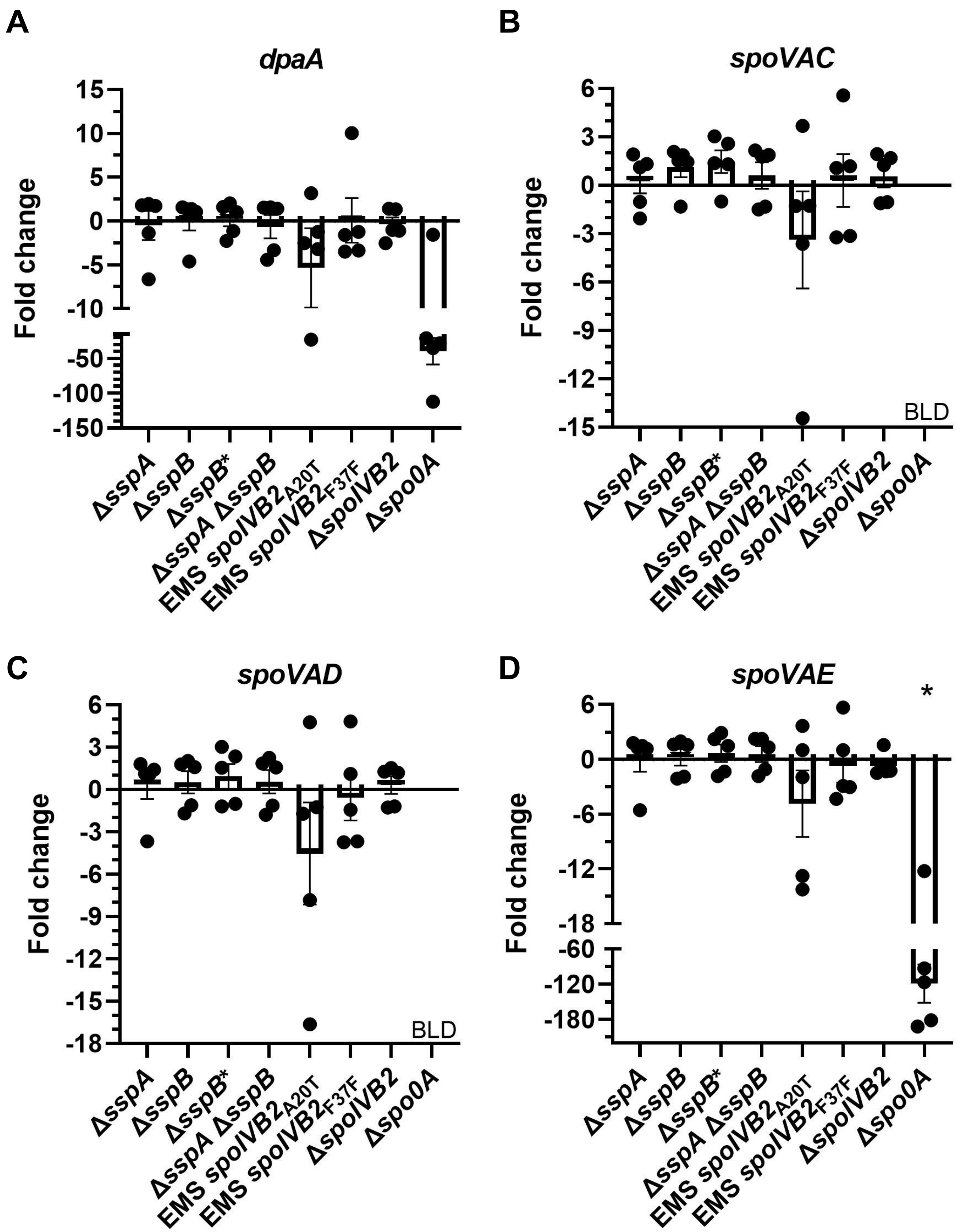

A
dpaA
Fold change
15
10
5
0
-5
-10
-50
-100
-150
B
spoVAC
Fold change
6
3
0
-3
-6
-9
-12
-15
BLD
C
spoVAD
Fold change
6
3
0
-3
-6
-9
-12
-15
-18
BLD
D
spoVAE
*
Fold change
6
0
-6
-12
-18
-60
-120
-180
ΔsspA
ΔsspB
ΔsspB*
ΔsspA ΔsspB
EMS spoIVB2A20T
EMS spoIVB2F37F
ΔspoIVB2
Δspo0A
